# The spike protein of SARS-CoV-2 induces endothelial inflammation through integrin α5β1 and NF-κB

**DOI:** 10.1101/2021.08.01.454605

**Authors:** Juan Pablo Robles, Magdalena Zamora, Gonzalo Martinez de la Escalera, Carmen Clapp

## Abstract

Vascular endothelial cells (EC) form a critical interface between blood and tissues that maintains whole-body homeostasis. In COVID-19, disruption of the EC barrier results in edema, vascular inflammation, and coagulation, the hallmarks of the severe disease. However, the mechanisms by which EC are dysregulated in COVID-19 are unclear. Here, we show that the spike protein of SARS-CoV-2 alone activates the EC inflammatory phenotype in a manner dependent on integrin α5β1 signaling. Incubation of human umbilical vein EC with whole spike, its receptor-binding domain, or the integrin-binding tripeptide RGD induced the nuclear translocation of NF-κB and enhanced the expression of leukocyte adhesion molecules VCAM1 and ICAM1, the adhesion of peripheral blood leukocytes, and the permeability of the monolayer. Inhibitors of integrin α5β1 activation prevented these effects. We suggest that the spike protein, through its RGD motif in the receptor-binding domain, binds to integrin α5β1 in EC to activate Rho GTPases, eNOS pathways, and the NF-κB gene expression program responsible for vascular leakage and leukocyte infiltration, respectively. These findings uncover a new direct action of SARS-CoV-2 on EC dysfunction and introduce integrin α5β1 as a promising target for treating vascular inflammation in COVID-19.

## Introduction

Endothelial cells (EC) dysfunction has emerged as a major driver of COVID-19^1–3^. During a resting state, EC maintain their barrier function by limiting vasopermeability and preventing coagulation and inflammation. However, when activated in response to damage or infection, EC produce chemoattractants, cytokines, and adhesion molecules, leading to vascular leakage, clot formation, inflammation, and leukocyte infiltration^4^. Most people with severe COVID-19 die from acute respiratory distress syndrome, pulmonary edema, cytokine storm, multiple organ failure, and disseminated intravascular coagulation^2^, all of which reflect EC dysfunction^1–3^. Moreover, severe cases or deaths due to COVID-19 associate with chronic endothelial damage from comorbidities such as aging, obesity, hypertension, diabetes, and cardiovascular disorders^5,6^.

Potential mechanisms of vascular dysfunction in COVID-19 include EC death in response to SARS-CoV-2 entry and replication^7^, the binding of the spike protein of SARS-CoV-2 to the ACE2 receptor causing its downregulation and subsequent mitochondrial dysfunction^8^, the activation of the proinflammatory kallikrein-bradykinin system, and the accumulation of proinflammatory and vasoconstrictor angiotensin II^2^. In addition, the spike protein can trigger the expression of proinflammatory cytokines and chemokines^9^, the production of toxic reactive oxygen species^10^, and cell death^11^. However, the underlying mechanisms of many of these effects remain unclear.

Alternative to ACE2, integrins may function as receptors mediating SARS-CoV-2 infection. Integrins are heterodimeric transmembrane cell adhesion molecules that exert various actions on hemostasis, inflammation, and angiogenesis. The spike protein contains an integrin-binding RGD motif exposed on the surface of the receptor-binding domain^12,13^ that binds to β1 integrins on pulmonary epithelial cells and monocytes^14^. In particular, blockage of the binding of the spike protein to the integrin α5β1 inhibits SARS-CoV-2 infection in vitro^11^ and in vivo^15^, demonstrating the therapeutic efficacy of targeting integrin α5β1 in COVID-19. Furthermore, fibronectin ligation of integrin α5β1 in EC activates the transcriptional factor NF-κB responsible for the expression of proteins involved in inflammation and angiogenesis^16^. These observations prompted us to investigate whether the binding of spike to the integrin α5β1 in EC is sufficient to induce the endothelium proinflammatory phenotype.

## Results

### Spike stimulates leukocyte adhesion to EC

Inflammatory changes can be assessed by the ability of EC to attach leukocytes, a hallmark of the inflammatory process. Peripheral blood leukocytes were incubated 1 h with human umbilical vein endothelial cells (HUVEC) pretreated for 16 h with spike, the receptor-binding domain of spike, the RGD tripeptide, or TNFα as a pro-inflammatory control (Figure 1). Spike stimulated the adhesion of leukocytes to HUVEC in a dose-dependent manner with high potency (EC_50_ = 1.6 nM), and this stimulation paralleled the one induced by TNFα, a main inducer of EC pro-inflammatory changes^17^ (Figure 1a, b). In addition, the receptor-binding domain of spike and the RGD tripeptide resulted in dose-response curves very similar to the one elicited by spike (EC_50_ = 1.8 nM) (Figure 1c). These results show that spike alone activates the proinflammatory program in EC and suggest that the RGD sequence located in the spike receptor-binding domain is responsible for this effect.

**Figure 1.**
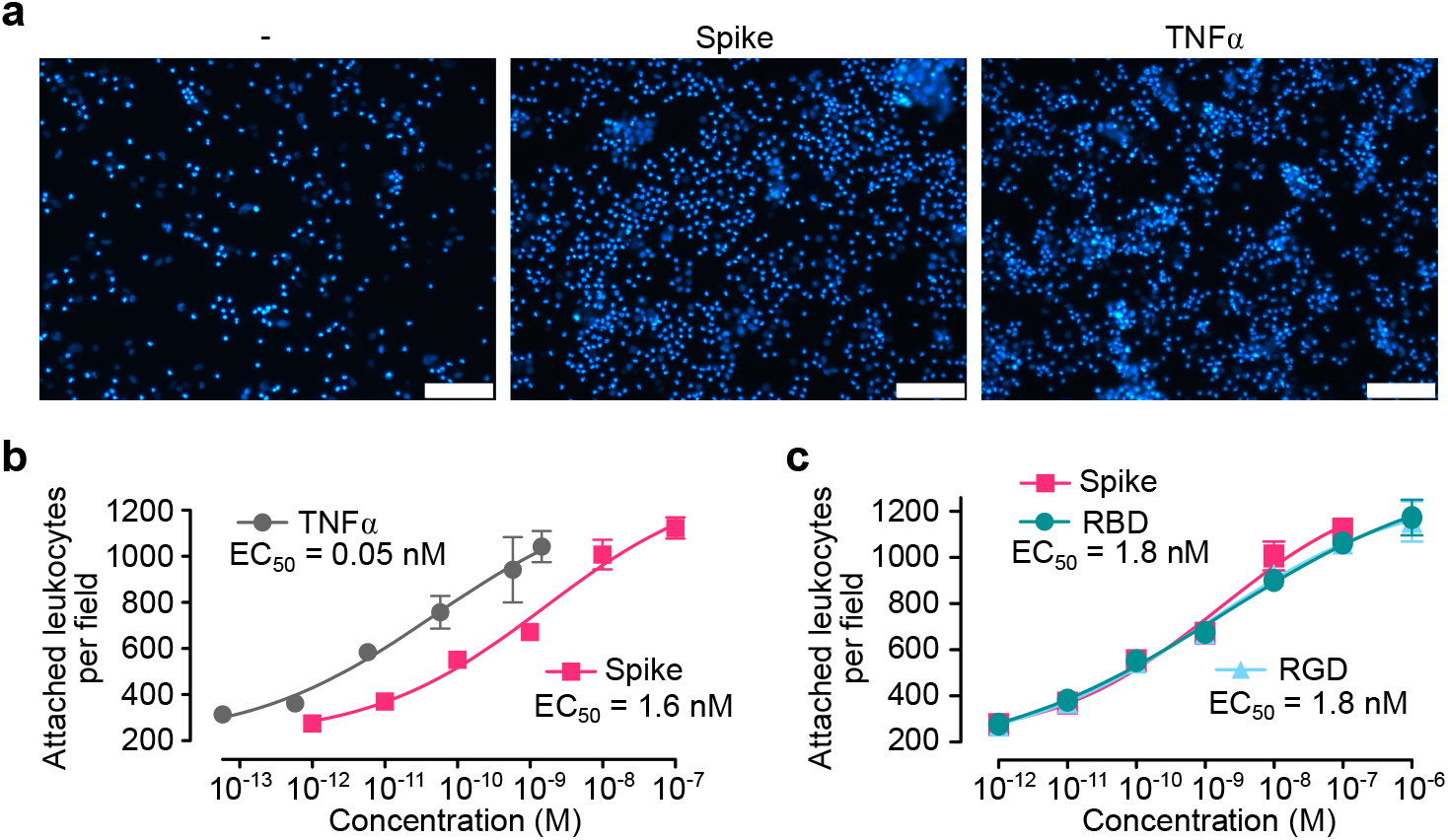
Spike-induced leukocyte adhesion to EC. **(a)** Micrographs showing leukocytes adhered to HUVEC monolayers incubated in the absence (-) or presence of spike (10 nM) or TNFα (1 nM). Scale bar = 120 μm. Dose-response stimulation of leukocyte adhesion to HUVEC in response to spike, TNFα **(b),** the receptor-binding domain of spike (RBD), or the RGD tripeptide **(c).** Values are means ± SD, *n* = 9. Dose-response curves were fitted by least square regression analysis (r^2^ > 0.93).

### Integrin α5β1 mediates spike-induced leukocyte adhesion to EC

RGD is the integrin-binding motif of several integrin ligands, including fibronectin, the main ligand of integrin α5β1. Fibronectin is upregulated during inflammation^18^ and stimulates the EC inflammatory program^16^. We used ELISA-based methods to confirm the binding of spike to integrin α5β1^11^. Plates were coated with integrin α5β1 and incubated with spike or the receptor-binding domain of spike in the absence or presence of the RGD tripeptide or neutralizing antibodies against integrin α5β1 or the α5 integrin subunit. Spike and its receptor-binding domain bound α5β1 integrin with the same affinity (Kd = 200 pM) (Figure 2a) and, as expected, the RGD tripeptide and the anti-α5β1 and -α5 antibodies prevented this binding (Figure 2b). Because fibronectin ligation of integrin α5β1 induces the expression of adhesion molecules in EC^16^, we explored whether the binding to integrin α5β1 mediated the spike-induced stimulation of leukocyte attachment. The effect of anti-α5β1 or anti-α5 antibodies was tested upon spike-, the spike receptor-binding domain-, the RGD tripeptide-, or TNFα-induced stimulation of leukocyte adhesion to HUVEC (Figure 2c). Consistent with an integrin α5β1-dependent effect, both antibodies blocked leukocyte adhesion in response to spike, the spike receptor-binding domain, and the RGD tripeptide. The effect of TNFα was not modified, confirming that it is integrin independent^19^.

**Figure 2.**
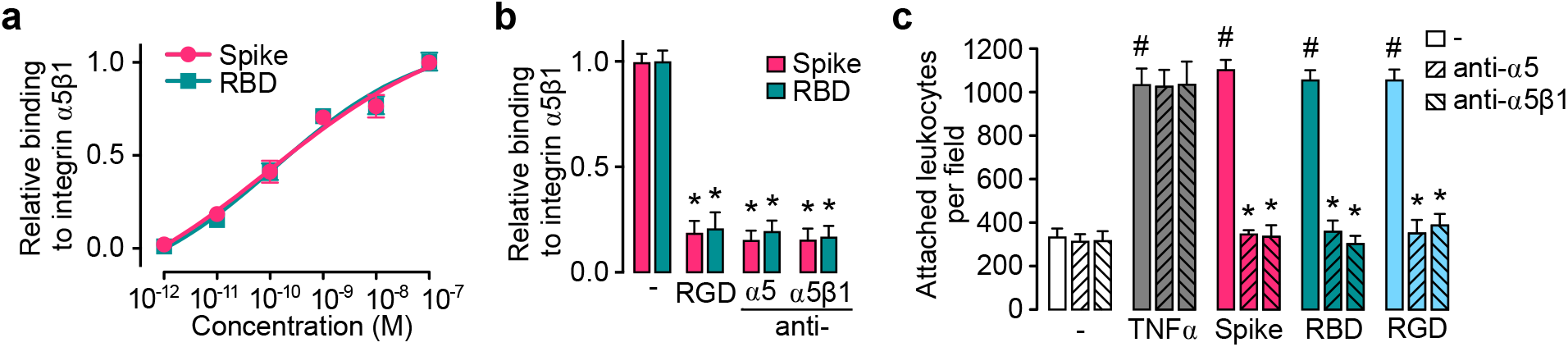
Integrin α5β1 mediates spike-induced stimulation of leukocyte adhesion to EC. **(a)** Binding of different doses of spike or spike receptor-binding domain (RBD) to immobilized integrin α5β1 in an ELISA-based assay. Binding curves were fitted by least square regression analysis (r^2^ > 0.97). **(b)** Effect of the RGD tripeptide, anti-integrin α5β1 antibodies or anti-α5 antibodies on the binding of spike or RBD to immobilized integrin α5β1. *P<0.001 versus basal binding in the absence of spike or RBD (-) (Two-way ANOVA, Šídák). **(c)** Leukocyte adhesion to a HUVEC monolayer in the absence (-) or presence of 100 nM spike, RBD, RGD, or 1nM TNFα, alone or together with antibodies against α5β1 integrin or the α5 integrin subunit. Values are means ± SD, *n* = 9, #P<0.001 versus (-), *P <0.001 versus absence of antibodies (Two-way ANOVA, Tukey).

### Spike-induced leukocyte adhesion to EC is dependent on integrin α5β1-mediated NF-κB activation

Because integrin α5β1 activates NF-κB in EC to elicit inflammation^16^, we asked whether the mechanism by which spike promotes leukocyte adhesion involved the activation of NF-κB. Inactive NF-κB is usually bound to a family of cytoplasmic inhibitory proteins, and its activation requires the degradation of these inhibitors followed by its nuclear translocation. The NF-κB cellular distribution was studied in HUVEC using fluorescence cytochemistry and a monoclonal antibody against the p65 subunit of NF-κB (Figure 3a). In the absence of treatment, p65 was homogeneously distributed throughout the cytoplasm of cells. Spike induced the accumulation of p65 in the cell nucleus, and this redistribution was like the one induced by TNFα. Anti-α5 antibodies blocked the spike-induced nuclear localization of p65, but not in response to TNFα; whereas the NF-κB inhibitor BAY11-7085 blocked the nuclear translocation of p65 induced by both spike and TNFα (Figure 3a). We conclude that spike activates NF-κB through its interaction with integrin α5β1.

**Figure 3.**
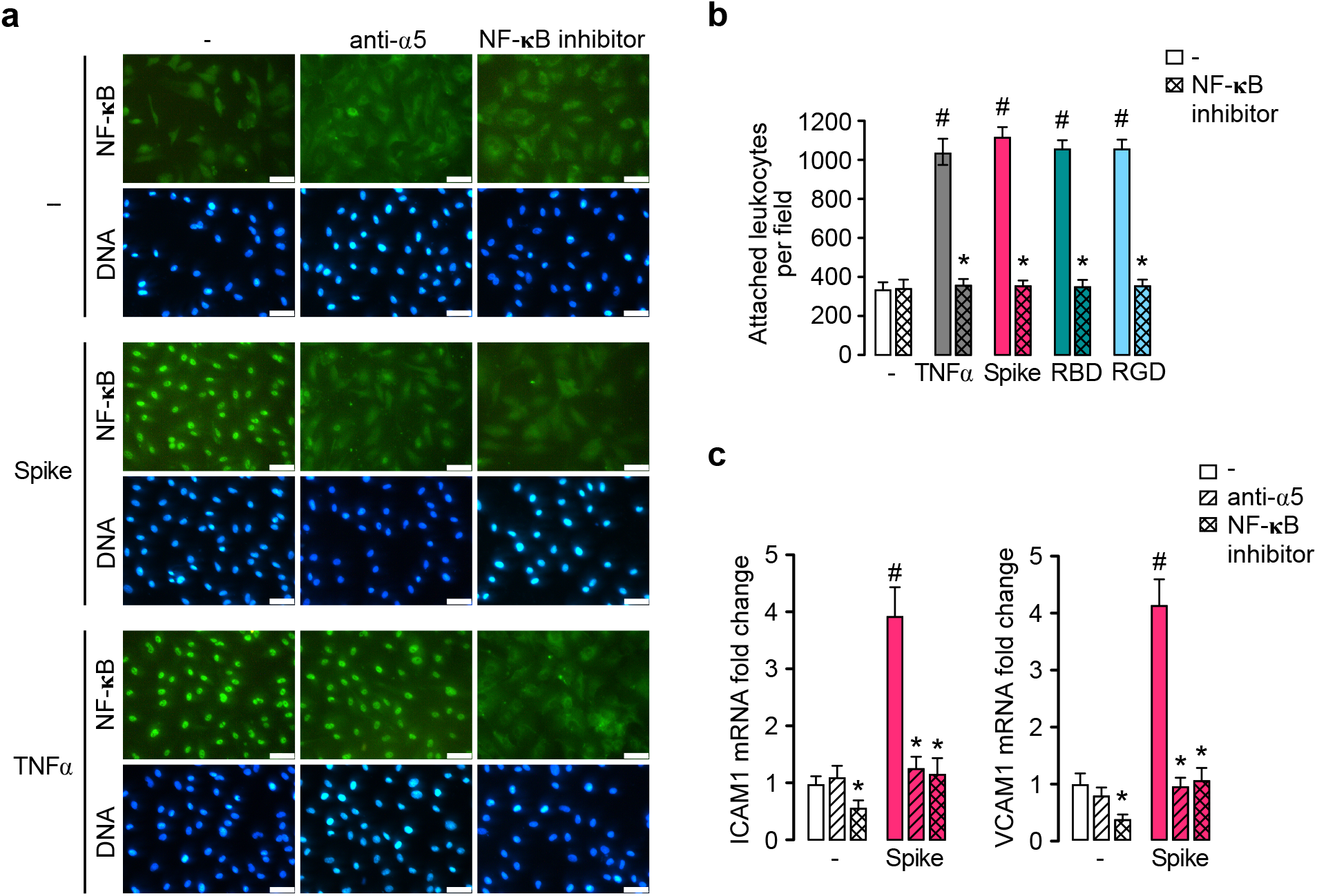
Spike-induced leukocyte adhesion to EC is dependent on integrin α5β1-mediated NF-κB activation. **(a)** Immunofluorescence detection of NF-κB (p65) in HUVEC incubated in the absence (-) or presence of spike (100 nM) or TNFα (1 nM) alone or in combination with anti-α5 antibodies or the NF-κB inhibitor, BAY 11-7085. Scale bar = 50 μm. **(b)** Leukocyte adhesion to HUVEC stimulated or not (-) by 1nM TNFα or 100 nM spike, spike receptor-binding domain (RBD), or the RGD tripeptide alone or together with BAY 11-7085. **(c)** Expression of intercellular adhesion molecule 1 (ICAM1) and vascular adhesion molecule 1 (VCAM1) in HUVEC incubated with or without (-) 100 nM spike, in the absence or presence of BAY 11-7085 or anti-α5 antibodies. Values are means ± SD, *n* = 9., #P < 0.001 versus the unstimulated control (-), *P < 0.001 versus absence of BAY 11-7085 or anti-α5 antibodies (Two-way ANOVA, Tukey).

To evaluate whether the spike-induced leukocyte adhesion to EC is dependent on NF-κB activation, we tested the effect of BAY11-7085 upon spike-, spike receptor-binding domain-, RGD tripeptide-, or TNFα-induced stimulation of leukocyte adhesion to HUVEC (Figure 3b). Inhibition of NF-κB prevented the stimulation of leukocyte adhesion in response to all treatments (Figure 3b). Furthermore, spike elevated by 4-fold the expression of intercellular adhesion molecule 1 (ICAM1) and vascular adhesion molecule 1 (VCAM1), both of mediate the firm adhesion of leukocytes to the apical surface of EC^20^; and the increased expression was prevented by the inhibition of NF-κB and integrin α5-immunoneutralization (Figure 3c). Altogether, our findings show that the binding of spike to integrin α5β1 activates the NF-κB pathway in EC responsible for leukocyte infiltration.

### Spike induces the hyperpermeability of EC monolayers via integrin α5β1

Besides leukocyte recruitment, β1 integrins promote vascular permeability, another critical aspect of inflammation^21^. Treatment with spike, the receptor-binding domain of spike, and the RGD tripeptide induced a drop in the trans-endothelial electrical resistance (TEER) of the HUVEC monolayer, indicative of hyperpermeability (Figure 4a). The drop was rapid, maximal at 30 minutes, and sustained thereafter. Furthermore, immunofluorescence showed the formation of actin stress fibers, EC retraction, and inter-endothelial gaps after 30 minutes of treatment with spike (Figure 4b). Likewise, spike interfered with the peripheral distribution of CD31, an adhesion protein that maintains the junctional integrity of EC^22^ (Figure 4b). These observations provided direct evidence of spike-induced increase in EC permeability.

**Figure 4.**
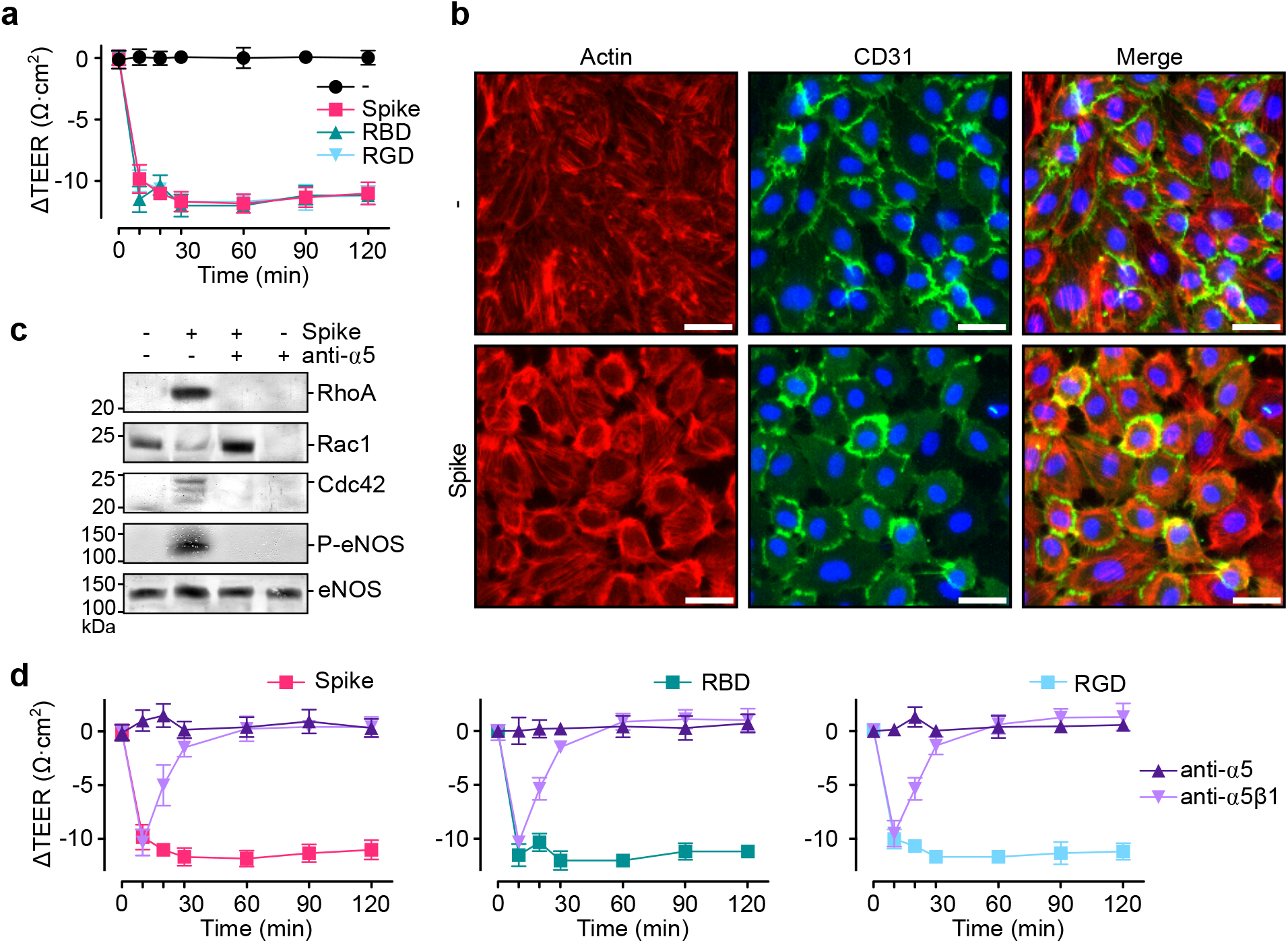
Spike induces the hyperpermeability of EC monolayers via integrin α5β1. **(a)** Changes throughout time in the transendothelial electrical resistance (ΔTEER) of HUVEC monolayers incubated without (-) or with 100 nM spike, spike receptor-binding domain (RBD), or RGD tripeptide. **(b)** Representative fluorescence cytochemistry of a HUVEC monolayer doubled-stained for F-actin and CD31 incubated without (-) or with 100 nM spike. Scale bar = 35 μm. **(c)** Representative western blot analysis of active RhoA, Rac1, Cdc42, phospho-eNOS (P-eNOS; relative to total eNOS) of lysates from HUVEC monolayers incubated without or with 100 nM spike in the absence or presence of anti-α5 antibodies. **d.** ΔTEER of HUVEC monolayers incubated throughout 120 minutes without or with 100 nM spike, RBD, or RGD in the presence or absence of antibodies against integrin α5β1 or the α5 integrin subunit. Values are means ± SD, *n* = 6.

RhoA, Rac1, and Cdc42 are Rho small guanosine triphosphatases (GTPases) that control the actin cytoskeleton and regulate EC barrier function^23^. Redistribution of actin stress fibers causing EC contraction involves RhoA activation^24^, whereas Rac1 and Cdc42 induce cell spreading^25^. Western blot analysis of HUVEC lysates incubated with spike for 30 min showed increases in RhoA and Cdc42 and a decrease in Rac1 that were blocked by the immunoneutralization of α5 (Figure 4c). Furthermore, endothelial nitric oxide synthase (eNOS) derived NO stimulates vasopermeability^26^, and spike promoted the phosphorylation/activation of eNOS in HUVEC in an α5-dependent manner (Figure 4c). Finally, immunoneutralization of integrin α5β1 prevented spike-, spike receptor-binding domain-, and RGD tripeptide-induced reduction in TEER (Figure 4d). Taken together, these studies show that spike binding to integrin α5β1 regulates Rho GTPases and eNOS phosphorylation to promote EC hyperpermeability.

### Volociximab and ATN-161 reduce spike-induced stimulation of EC leukocyte adhesion and permeability

Inhibitors of integrin α5β1 have been developed as promising therapeutics. For example, volociximab, a chimeric anti-integrin α5β1 monoclonal antibody, is under clinical evaluation for the treatment of cancer^27^; and preclinical studies have evaluated the efficacy of the integrin α5β1 binding peptide, ATN-161, to inhibit beta-coronavirus^28^ and SARS-CoV-2 virus infections^11,15^. Here, we show that both volociximab and ATN-161 block the binding of spike and the spike receptor-binding domain to α5β1immovilized in ELISA plates (Figure 5a); and prevent the leukocyte adhesion to HUVEC (Figure 5b) and HUVEC hyperpermeability (Figure 5c) in response to spike, spike receptor-binding domain, and RGD tripeptide.

**Figure 5.**
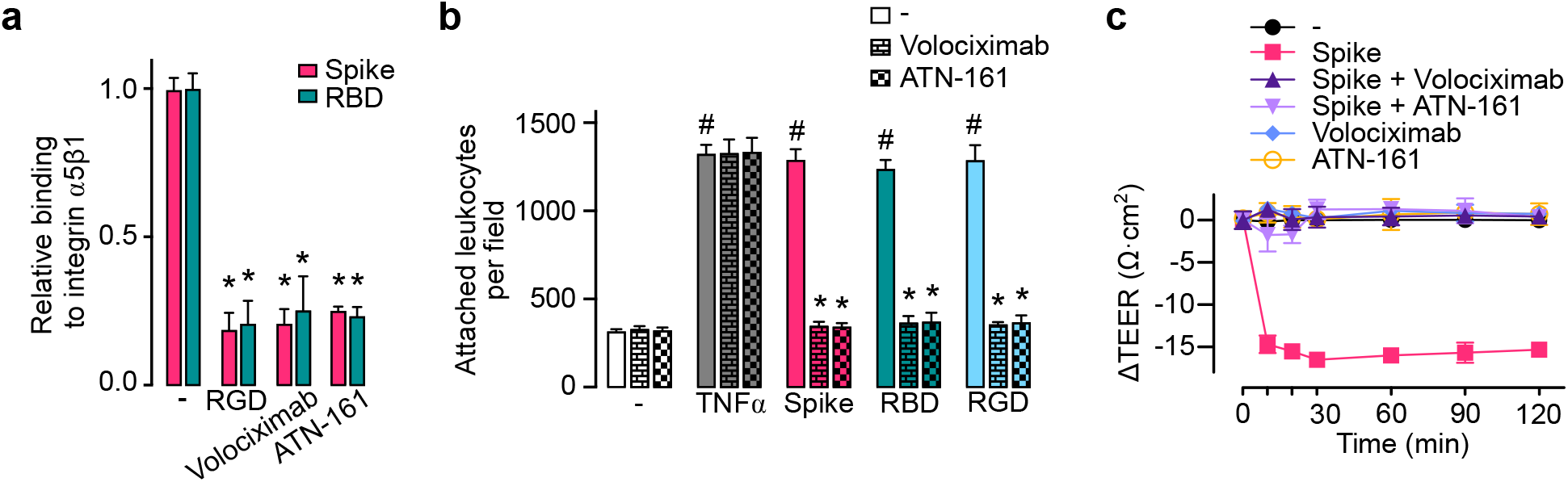
Volociximab and ATN-161 reduce spike-induced stimulation of EC leukocyte adhesion and permeability. **(a)** Binding of 100 nM spike or spike receptor-binding domain (RBD) to immobilized integrin α5β1 in the absence (-) or presence of the RGD tripeptide or the integrin α5β1 inhibitors, volociximab (5 μg mL^−1^) and ATN-161 (500 nM). **(b)** Leukocyte adhesion to HUVEC incubated without (-) or with 1 nM TNFα or 100 nM spike, RBD, or RGD, in the absence or presence of volociximab (5 μg mL^−1^) or ATN-161 (500 nM). #P < 0.001 versus the unstimulated control (-), *P < 0.001 versus absence of integrin α5β1 inhibitors (Two-way ANOVA, Tukey). **(c)** Changes throughout time of the transendothelial electrical resistance (ΔTEER) of HUVEC monolayers incubated without (-) or with 100 nM spike in the presence or absence of volociximab (5 μg mL^−1^) or ATN-161 (500 nM). ΔTEER values of HUVEC incubated only with volociximab or ATN-161 are also shown. Values are means ± SD, *n*= 6.

## Discussion

Accumulating evidence has defined COVID-19 as a vascular disease^1–3,29^. Blood vessel injury causes progressive lung damage and multi-organ failure in severe COVID-19, owing to edema, intravascular coagulation, vascular inflammation, and deregulated inflammatory cell infiltration. Multiple mechanisms have been proposed for vascular dysfunction in COVID-19^1,2,29^; however, little is known regarding the direct action of SARS-CoV-2 on EC^10,30^.

ACE2 is the best-established host receptor for spike^31–33^, although other spike cell surface receptors have been described, i.e., neuropilin-1^34^, toll-like receptos^9,35^, and RGD-binding integrins^11,14^. In particular, integrin α5β1 is an RGD-binding integrin that, upon spike binding, mediates SARS-CoV-2 entry and infection of epithelial cells and monocytes in vitro^11^ and increases lung viral load and inflammation in vivo^15^. Ligation of integrin α5β1 by fibronectin RGD motif activates the expression of proinflammatory genes in EC^16^, but the binding of spike to α5β1 in EC and its impact on the EC inflammatory response has not been addressed.

In this work, we show that spike binding to integrin α5β1 activates the inflammatory program of EC. Spike stimulated the expression of adhesion molecules ICAM1 and VCAM1 and the attachment of leukocytes to EC monolayers like TNFα, a well-known inducer of sustained EC inflammatory responses^17^. The fact that both the RGD tripeptide and the receptor-binding domain of spike elicited almost identical responses to those of spike, that the RGD tripeptide itself blocked spike- and spike receptor-binding domain-binding to α5β1, and that α5β1-neutralizing antibodies prevented the spike-induced proinflammatory effect in EC indicated that spike, through its RGD motif in its receptor-binding domain, binds to integrin α5β1 in EC to promote inflammation. Furthermore, we show that the transcription factor NF-κB is a primary contributor to α5β1 signaling in EC in response to spike. Spike and its receptor-binding domain stimulated NF-κB nuclear translocation, and inhibitors of NF-kB and α5β1 prevented the spike-induced expression of ICAM1 and VCAM1 and leukocyte adhesion to EC. These findings are consistent with a previous study showing that NF-κB is the signaling pathway by which fibronectin ligation to α5β1 upregulates proinflammatory genes in EC^16^. Also, NF-κB is a predominant signaling molecule activated in EC by proinflammatory cytokines, such as TNFα via an integrin-independent route^17^.

Neutrophils, macrophages, and lung epithelial cells react to spike^9,35^ and to the envelope protein^36^ via the activation of toll-like receptor-induced cytokine production, a mechanism underlying the cytokine storm observed in severe COVID-19^37^. Integrin α5β1 is widely expressed in immune cells^16^, and it may cooperate with toll-like receptors to boost their signaling pathways^38^. However, human EC express few or no toll-like receptors on their surface^39^, implying that interactions between toll receptors and integrin α5β1 do not contribute to the proinflammatory effect of spike in the endothelium.

Excessive vasopermeability leading to edema is another hallmark of severe COVID-19. Here, we showed that spike binding to integrin α5β1, through its RGD motif, stimulated the hyperpermeability of EC monolayers. Spike-induced hyperpermeability was mimicked by its receptor-binding domain and the RGD tripeptide and prevented by neutralizing antibodies against the α5β1 integrin and α5-integrin subunit.

β1-integrins are known to stabilize EC-cell junctions during development^40^, but also to promote their disruption under the context of inflammation^21^. Several inflammatory agents (LPS, IL-1β, and thrombin) signal via β1-integrin to promote EC permeability and contractility in vitro and vascular leakage in vivo^41^. Consistent with this observation, we showed that the spike signals through α5β1to promote the formation of stress fibers, endothelial cell retraction, and inter-endothelial gap formation. These processes are under the control of GTPases of the Rho family (RhoA, Rac, Cdc42), and spike upregulated RhoA and Cdc42 and downregulated Rac1 in EC. Rho GTPases signal downstream of integrin binding to exert both positive and negative effects on EC junctions and EC matrix adhesion depending on its context. Under inflammation, RhoA associates with loss of barrier integrity, Rac1 with maintenance, and Cdc42 with barrier stabilization and recovery^42^.

Consistent with the above findings, spike interfered with the peripheral distribution of CD31 observed in stable EC monolayers. Because CD31 exhibits adhesive properties and is concentrated mainly at junctions between adjacent cells^22^, its altered localization in EC is consistent with increased vasopermeability. This observation agrees with a recent report showing that spike downregulates the expression of EC junctional proteins from normal and diabetic mice, reflecting the disruption of the endothelial barrier integrity^43^. Finally, spike stimulated the phosphorylation/activation of eNOS in EC, and eNOS-derived NO is a principal vasorelaxant and vasopermeability factor^26,44^.

Altogether, we have shown that, by binding to α5β1, spike changes the EC phenotype to promote vascular inflammation. Upon spike-mediated α5β1 activation, EC lose their ability to control permeability or quiesce leukocytes, both of which are characteristics of EC dysfunction^17^. These findings provide insights into the mechanisms making COVID-19 a vascular disease and encourage the development of therapeutic approaches that directly focus on α5β1-mediated vascular changes.

Finally, we showed that two inhibitors of α5β1, volociximab and ATN-161, developed as promising therapeutics, blocked spike-induced leukocyte adhesion and hyperpermeability of EC. Indeed, ATN-161 has already been shown to inhibit SARS-CoV-2 virus infection in vivo^15^. Our findings, add EC as a target for these drugs to reduce vascular inflammation in COVID-19, suggest the therapeutic potential of agents commonly used to treat vascular diseases, and provide tools for guiding research into the molecular mechanisms mediating the pathophysiology of COVID-19.

## Methods

### Reagents

Recombinant SARS-CoV-2-His-Tagged spike (Active Trimer) (cat. 10549-CV) was purchased from Bio-techne (Minneapolis, MI, USA). Recombinant receptor-binding domain of spike (amino acids 319-541) flanked by the signal peptide (amino acids 1-14) and containing a 6xHis tag was produced as previously reported^45^. Arg-Gly-Asp (RGD) tripeptide, the NF-κB activation inhibitor BAY 11-7085, and ATN-161 were purchased from Sigma Aldrich (St. Louis, MO, USA), and human recombinant TNFα was from R&D Systems (Minneapolis, MN, USA). Purified human integrin α5β1 and anti-α5 (MAB1956Z) and anti-α5β1 (MAB1999) antibodies were from Merck (Kenilworth, NJ, USA). Volociximab was from Novus Biologicals (Littleton, CO, USA).

### Cell culture

Human umbilical vein endothelial cells (HUVEC) were obtained as described^46^. HUVEC were maintained in F12K media supplemented with 20% fetal bovine serum (FBS), 100 μg mL^−1^ heparin (Sigma-Aldrich), 25 μg mL^−1^ endothelial cell growth supplement (ECGS, Corning, NY, USA), and 100 U mL^−1^ penicillin-streptomycin.

### Leukocyte adhesion assay

HUVEC were seeded on a 96 well plate and grown to confluency. HUVEC monolayers were treated for 16 hours with TNFα, spike, the spike receptor-binding domain, and the RGD tripeptide alone or in combination with anti-α5β1 antibodies (5 μg mL^−1^), anti-α5 antibodies (5 μg mL^−1^), volociximab (5 μg mL^−1^), or ATN-161 (500 nM) in 20% FBS F12K without heparin or ECGS. For inhibition of NF-κB, cells were pretreated for 30 minutes with 5 μM of BAY 11-7085. Leukocytes were prepared from whole blood collected into heparin tubes. Briefly, blood was centrifuged (300 x g for 5 minutes) and the plasma layer discarded. The remaining cell pack (~5 mL) was diluted in 45 mL of red blood cells lysis buffer containing150 mM of NH_4_Cl, 10 mM NaHCO_3_, and 1.3 mM EDTA (disodium) and gently rotated for 10 minutes at RT. Leukocytes were collected by centrifugation (300 x g for 5 min), the pellet washed with cold PBS followed by centrifugation (300 x g for 5 min), and resuspended into 5 mL of warm PBS containing 5 μg mL^−1^ Hoechst 33342 (Thermo Fisher Scientific, Waltham, MA, USA) to stain live leukocytes.

Leukocytes were incubated in darkness for 30 minutes, washed with PBS three times by centrifugation (200 x g for 5 min each), resuspended into supplemented culture media, and taken to 10^6^ cells mL^−1^. After 16 hours of treatment, the medium of HUVEC monolayers was replaced with 100 μL of medium containing Hoechst-stained leukocytes (~100,000 leukocytes per well) and cells incubated for 1 hour at 37°C. HUVEC monolayers were washed three times with warm PBS, and images were obtained in an inverted fluorescent microscope (Olympus IX51, Japan) and quantified using the CellProfiler software^47^.

### ELISA

A 96-well ELISA microplate was coated overnight at 4 °C with 25 ng per well of human integrin α5β1 diluted in PBS, and blocked for 1 hour at RT with 5% w/v nonfat dry milk in 0.1% Tween-20-PBS (PBST). After blocking, microplates were washed three times with PBST and different concentrations of spike or spike receptor-binding domain added. Anti-α5β1 antibodies (5 μg mL^−1^), the RGD tripeptide (500 nM), ATN-161 (500 nM), or volociximab (5 μg mL^−1^) were added together with 100 nM of spike or spike receptor-binding domain. Dilutions were in 0.2 mg mL^−1^ BSA-PBST. Microplates were then incubated for 1 hour at RT, washed three times with PBST, incubated for 1 hour at RT with 1:1,000 of anti-spike antibody (Rabbit MAb, Sino Biological, Beijing, CN) diluted in blocking buffer, washed (three times in PBST), and incubated (1 hour at RT) with horseradish peroxidase-labeled goat anti-rabbit secondary antibodies (Thermo Fisher Scientific) diluted 1:2,500 in 50% blocking buffer. After a three-wash step, the microplates were incubated with 100 μL of an OPD substrate tablet diluted in 0.03% H_2_O_2_ citrate buffer for 30 minutes in darkness. The reaction was stopped with 50 μL of 3M HCl, and absorbance was measured at 490 nm.

### NF-κB nuclear translocation analysis

HUVEC were seeded on 18 mm-coverslips coated with fibronectin (1 μg cm^−1^) and placed in a 12-well plate. Cells were grown in complete media to 80% confluence and, on the day of the assay, the medium was replaced with 0.5% FBS-F12K. The NF-κB activation inhibitor, BAY 11-7085 (5 μM), or anti-α5 antibodies (5 μg mL^−1^) were added 30 minutes prior to a 30-minutes incubation with spike (100 nM) or TNFα (1 nM). Cells were washed, fixed in 4% PFA for 30 minutes at RT, permeabilized for 30 minutes with 0.5% Tx-100-PBS, blocked with 0.05% Tx-100, 1% BSA, 5% normal goat serum-PBS at RT for 1 hour, and incubated overnight at 4°C in a humid chamber with 1:200 mouse monoclonal anti-NF-κB p65 antibody (Santa Cruz, CA, USA) in 0.1% Tx-100, 1% BSA-PBS. Cells were washed and incubated under darkness at RT with a goat anti-mouse secondary antibody (1:500, Alexa fluor 488, Abcam, Cambridge, UK) in 0.1% Tx-100, 1% BSA-PBS for 2 hours. Nuclear DNA was counterstained with 5 μg mL^−1^ Hoechst 33342 (Sigma-Aldrich) and the coverslips washed and mounted with Vectashield mounting medium (Vector laboratories, Burlingame, CA) and observed under fluorescence microscopy (Olympus IX51).

### qPCR

HUVEC grown to 80% confluency on 6-well plates were incubated under starving conditions (0.5% FBS, F12K medium) for 30 minutes with the NF-κB activation inhibitor, BAY 11-7085 (5 μM) or anti-α5 antibodies (5 μg mL^−1^) followed by a 4-hour incubation with 100 nM spike. RNA was isolated using TRIzol reagent (Invitrogen, Carlsbad, CA, USA), retrotranscribed with the High-Capacity cDNA Reverse Transcription kit (Applied Biosystems, Foster City, CA, USA), and quantified using Maxima SYBR Green qPCR Master Mix (Thermo Fisher Scientific) in a final reaction of 10 μL containing 20 ng of cDNA, and 0.5 μM of each of the following human primers: VCAM1 forward (5’-gcactgggttgactttcagg-3’) and reverse (5’-aacatctccgtaccatgcca-3’); ICAM1 forward (5’-gtgaccgtgaatgtgctctc-3’) and reverse (5’-cctgcagtgcccattatgac-3’); and GAPDH forward (5’-gaaggtcggagtcaacggatt-3’) and reverse (5’-tgacggtgccatggaatttg-3’).

### Vasopermeability assay

HUVEC were grown to confluence on a 6.5 mm transwell with a 0.4 μm pore coated with 1 μg cm^−1^ fibronectin (Thermo Fisher Scientific). Trans-endothelial electrical resistance (TEER) was measured using the epithelial EVOM^2^ Volt/Ohm meter (World Precision Instruments, Sarasota, FL, USA). The assay was performed once the TEER measurement was stable for at least two days. Cells were pretreated for 5 minutes with anti-α5 antibodies (5 μg mL^−1^), anti-α5β1 antibodies (5 μg mL^−1^), volociximab (5 μg mL^−1^), or 500 nM ATN-161 followed by the treatment with 100 nM spike, spike receptor-binding domain, or the RGD tripeptide. TEER measurements were made over 120 minutes.

### Fluorescence cytochemistry for CD31 and F-Actin

HUVEC were seeded on 18 mm coverslips coated with fibronectin (1 μg cm^−1^) and placed in a 12-well plate. Cells were grown to confluence and, once the monolayer was completely formed, cells were starved (0.5% FBS) for 1 hour before the addition of 100 nM of spike for 30 minutes. Cells were washed, fixed with 4% PFA for 30 minutes, permeabilized with 0.5% Tx-100-PBS for 30 minutes, blocked with 0.1% Tx-100, 1% BSA, 5% normal goat serum-PBS for 1 hour at RT, and incubated overnight at 4°C in a humid chamber with 1: 100 anti-CD31 antibody (Abcam) in 0.1% Tx-100, 1% BSA-PBS. Cells were washed and incubated in darkness with goat anti-mouse secondary antibodies (1:500, Alexa fluor 488, Abcam, Cambridge, UK) in 0.1% Tx-100, 1% BSA-PBS for 2 hours. Cells were then washed with PBS and 300 μl of 160 nM rhodamine-phalloidin (Thermo Fisher Scientific) added to each well for 1 hour in darkness, followed by a PBS wash and nuclear DNA counterstaining with 5 μg mL^−1^ Hoechst 33342 (Sigma-Aldrich). Finally, the coverslips were washed, mounted, and observed under fluorescence microscopy.

### RhoA/Rac1/Cdc42 analysis

HUVEC were seeded on a 6-well plate and grown in complete media. Eighty-percent confluent cells were then starved for 3 hours (0.5% FBS) and treated or not for 30 minutes with the anti-α5 antibodies (5 μg mL^−1^) followed by the addition of 100 nM spike for a 30 minute-incubation. Cells were washed twice with ice-cold PBS, and the RhoA/Rac1/Cdc42 activation was evaluated using the RhoA/Rac1/Cdc42 combo activation assay kit (Abcam) according to the manufacturer’s instructions. The kit is based on the binding of the active form of RhoA to the Rho-binding domain of Rhotekin and of active Rac1 or Cdc42 to the p21-binding domain of p21 activated protein kinase (PAK1). Both, Rhotekin and PAK1 binding domains are coupled to agarose beads, and the isolation and detection of the active forms of RhoA, Rac1, and Cdc42 were done under conventional bead-precipitation and western blot protocols using goat anti-mouse alkaline phosphatase secondary antibodies (Jackson ImmunoResearch, Philadelphia, PA, USA, 1:5000) and a colorimetric detection kit (BioRad, Hercules, CA, USA).

### Western blot analysis of eNOS phosphorylation at Ser1177

HUVEC were seeded on a 6-well plate and grown in complete media to reach an 80% confluency. On the day of the assay, HUVEC were starved (0.5% FBS) for 3 hours and treated or not for 30 minutes with the anti-α5 antibody (5 μg mL^−1^) followed by the addition of 100 nM spike for a 30 minute-incubation. Cells were washed twice with cold TBS, scraped with 200 μL RIPA buffer supplemented with 1:100 halt protease-phosphatase inhibitor cocktail (Thermo Fisher Scientific) and 5 mM of EDTA, centrifuged (10,000 x g, 4°C for 10 min), and supernatants aliquoted and stored (−70°C) for western blot analysis. Forty-five μg of protein were resolved in SDS-PAGE and blotted with anti-phospho-eNOS (Ser1177) antibodies (Cell Signaling, Danvers, MA, USA, 1:250), followed by incubation with goat anti-rabbit horseradish peroxidase secondary antibodies (Jackson ImmunoResearch, 1:5000). Immunoblots were developed using the SuperSignal West Pico PLUS chemiluminescent substrate kit (Thermo Fisher Scientific) and the FluorChem E imager and gel documenter system (ProteinSimple, San Jose, CA, USA). Membranes were reblotted using antibodies against total eNOS (Cell Signaling, 1:500), goat anti-rabbit alkaline phosphatase secondary antibodies (Jackson ImmunoResearch, 1:5000), and a colorimetric detection kit (BioRad).

## Author Contributions

JPR, MZ, and CC conceived research, designed and supervised the study, interpreted the data, and wrote the manuscript. MZ performed most research. GM de la E revised the study. All authors reviewed and approved the manuscript.

## Availability of data and material

All data generated or analyzed during this study are included in this manuscript.

## Ethics approval

Not applicable.

## Acknowledgments

We thank Xarubet Ruíz Herrera, Fernando López Barrera, and Adriana González Gallardo for their excellent technical assistance.

